# Multisensory information about changing object properties can be used to quickly correct predictive force scaling for object lifting

**DOI:** 10.1101/2022.01.26.477825

**Authors:** Vonne van Polanen

## Abstract

Sensory information about object properties, such as size or material, can be used to make an estimate of object weight and to generate an accurate motor plan to lift the object. When object properties change, the motor plan needs to be corrected based on the new information. The current study investigated whether such corrections could be made quickly, after the movement was initiated. Participants had to grasp and lift objects of different weights that could be indicated with different cues. During the reaching phase, the cue could change to indicate a different weight and participants had to quickly adjust their planned forces in order to lift the object skilfully. The object weight was cued with different object sizes (Experiment 1) or materials (Experiment 2) and the cue was presented in different sensory modality conditions: visually, haptically or both (visuohaptic). Results showed that participants could adjust their planned forces based on both size and material and in all sensory modality conditions. Furthermore, the findings indicate that visual and haptic information were integrated in the visuohaptic condition, although the multisensory condition did not outperform the conditions with one sensory modality. These results suggest that motor plans can be quickly corrected based on sensory information about object properties from different sensory modalities. These findings provide insights into the information that can be shared between brain areas for the online control of hand-object interactions.

## 1 INTRODUCTION

Sensory information is essential to skilfully control movements. This information can be used for motor planning, e.g. predictive scaling of fingertip forces to an object’s weight in lifting movements. This predictive force scaling results in smooth lifting behaviour, since feedback processes are often too slow to allow for skilled movements (Johansson and Westling 1988). Since object weight cannot be directly perceived before contacting the object, other object properties, such as the size or material, can be used to estimate the object’s weight and generate an appropriate motor plan (Gordon et al. 1991b; Buckingham et al. 2009). When the weight prediction proves to be incorrect, online corrective feedback loops, likely relying on tactile afferents (Johansson and Flanagan 2009), can quickly adjust the motor plan (Johansson and Westling 1988), e.g. by increasing the forces when the object is heavier than expected.

When the expected properties of the object are incorrect, for example, when the object is larger than initially expected, a new weight estimation has to be inferred from memory (e.g. larger objects are heavier) to correct the motor plan. If such a correction is needed when the movement has already been initiated, this poses time constraints on the correction processes. Previously it has been argued that such online corrections could not be performed quickly, since they would rely on slow memory processes. Specifically, the dual-stream theory (Goodale and Milner 1992; Milner and Goodale 2008) suggests a distinction between a dorsal stream, running from the occipital cortex to the parietal cortex, for the online control of movements, and a ventral stream (occipital to temporal cortices) concerned with object recognition and weight information (Gallivan et al. 2014). Originally, the theory suggested that the separate dorsal stream would not have quick access to the ventral stream information relating to memorized object properties and quick corrections would hence not be possible, because corrections would have to rely on slower processes from the ventral stream.

Current views acknowledge interactions between the two streams (Cloutman 2013; van Polanen and Davare 2015; Milner 2017), but the time scale of sharing information is still rarely investigated. Importantly, it has been shown that fast online corrections in object lifting based on visual cues are in fact possible, based on a visual change in object size after initiation of the reaching movement (Brouwer et al. (2006)) or a change in a colour cue at the go-signal (van Nuenen et al. 2012). These studies indicate that visual cues about object properties that indicate object weight can be used to perform relatively quick online corrections in force scaling. However, it remains unclear whether corrections can also be made based on other object properties besides size and colour. This is especially of interest since different object properties are generally processed at different locations in the brain (e.g. size and texture, (James et al. 2007; Cant et al. 2009) and might not be equally available for online control.

In addition, the previously mentioned studies (Brouwer et al. 2006; van Nuenen et al. 2012) only examined corrections based on visual information, whereas haptic information about object properties can also be used for force scaling (Gordon et al. 1991a). Similar to the visual system, a division between the processing for online control and object perception has been suggested for the somatosensory system (Dijkerman and de Haan 2007). In practice, haptic information is often available from one hand holding the object while the other hand is reaching towards this object. In this case, sensory information also needs to be transferred between hemispheres, which might take more time. Therefore, it is of interest to study whether haptic information about object properties can be used for fast online control of hand movements.

When both visual and haptic information are available, the multisensory information can be integrated. Studies on object perception, where the accuracy of a perceptual estimation is investigated, showed that sensory inputs are often integrated optimally (Ernst and Banks 2002). That is, the multisensory estimate is based on an optimal weighting of the unimodal inputs based on their accuracy, which improves performance compared to the unimodal perceptual estimates alone. In contrast to research on perceptual judgements, less is known about multisensory integration for online control of movements, especially when handling objects. Because of time constraints in online control, not only the accuracy but also the delay of information is important (Crevecoeur et al. 2016). It is known that inputs from multiple senses are used for hand shaping (see for a recent review: Betti et al. 2021) and object lifting (van Polanen et al. 2019) and can even improve object grasping (Camponogara and Volcic 2019b). Less is clear about the specific weighting of visual and haptic information. Some grasping studies suggest that is mostly relied on vision (Pettypiece et al. 2010), while faster corrections to grasping movements were found for haptic information (Camponogara and Volcic 2019a). Since the weighting can be adjusted to tool properties when handling objects with tools (Takemura et al. 2016), it is possible that multisensory integration is altered in the case of corrective movements.

The aim of the present study was to investigate how corrections in force scaling can be made in response to changes in sensory information about the object to be lifted. Participants grasped and lifted objects of different weights, which was indicated by a cue before they started their movement in order to plan their forces. Importantly, sometimes the cue was changed after they initiated their reaching movement so they had to adjust their motor plan to lift the object skilfully. In this way, it could be examined whether participants could correct their motor plan online within a short timescale based on sensory cues.

The type and sensory modality of the cues were altered to address two questions. First, cues were either presented visually, haptically or visuohaptically to be able to examine the role of sensory modality and multisensory integration. Second, in one experiment the object weight was indicated with object size, whereas in the second experiment the weight was indicated with material cues. Since object processing is organised differently in the brain for shape and texture, indicative of material, (James et al. 2007; Cant et al. 2009), this information might also be differently available for online corrections. The hypotheses are that participants are able to correct their motor plan based on sensory information, irrespective of the type or sensory modality of the cue. Furthermore, it was expected that corrective performance would be improved in the multisensory condition compared to the unimodal conditions.

## 2 METHODS

### 2.1 Participants

Thirty right-handed participants were recruited. Fifteen participants took part in Experiment 1 (11 females, 24.3±3.2 years) and another 15 participated in Experiment 2 (12 females, 23.8±3.5 years). Handedness was assessed with the Edinburgh handedness questionnaire (Oldfield 1971) and this gave an average LQ of 90±14 and 90±18, for Experiment 1 and 2, respectively. One additional participant only finished one of two sessions of Experiment 1 and was excluded from data analysis. All participants signed informed consent before performing the experiment. The experiments were approved by the local ethical committee of KU Leuven and were in accordance with the Declaration of Helsinki.

### 2.2 Apparatus

Two force-torque sensors (Nano17, ATI Industrial Automation) were used to measure force scaling (1000 Hz). The sensors were mounted on a manipulandum, which was attached to a carbon-fiber basket in which different objects could be placed (Fig. 1). The surface of the sensors, where the participants placed their fingertips, were covered with sandpaper (P600) to increase friction.

**Fig. 1.**
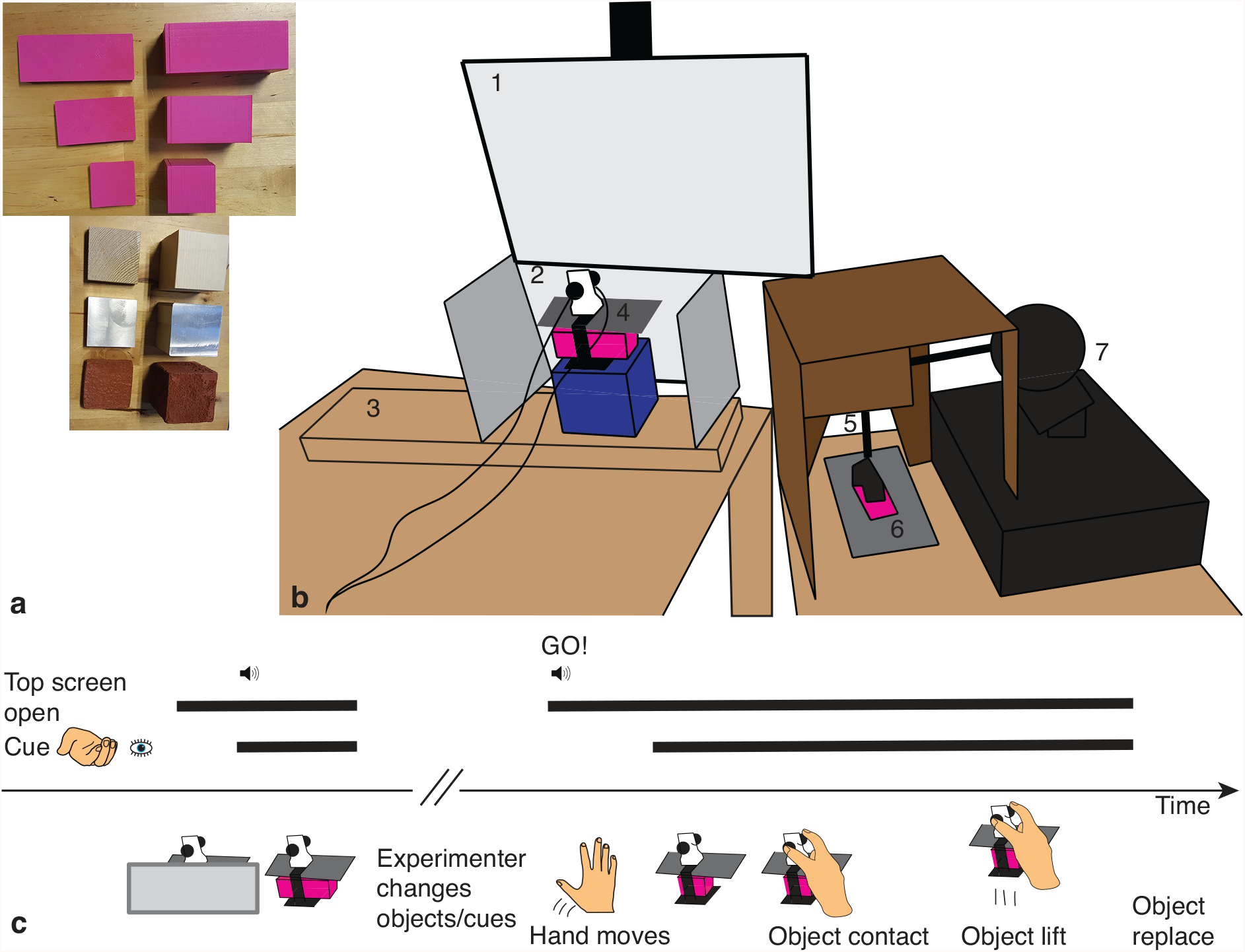
**a**. Haptic cues (left) and objects (right). Size cues were used in Experiment 1 (large, medium and small; top pink objects) and material cues in Experiment 2 (wood, metal and stone; bottom). **b**. Experimental setup. 1) Upper screen for showing force sensors. 2) Lower screen for showing object in manipulandum (4). The screen had two side panels to hold it in place and participants moved their hand over the panel. 3) Start position of the right hand. 4) The manipulandum, placed on a platform (blue box), with two force sensors, in which an object (pink cuboid) could be placed. The object was hidden under a paper cover (dark grey). 5) Opening in a cardboard box through which participants placed their left hand. 6) The left hand was placed on a cushion. The haptic cue (pink plate) was attached to the arm of a haptic device (7) and pressed on the hand palm. **c**. Experimental time line. The top screen opens to allow view of the sensors. The first cue is presented (visually, haptically or visuohaptically), accompanied by a beep. Next, the experimenter changes the objects and cues in case of a change condition. After the GO-signal, the hand starts to move and when it reaches a threshold height, the 2^nd^ cue is presented. The object is grasped, lifted and replaced

Hand position was measured (240 Hz) with an electromagnetic tracking system (trakSTAR, NDI) with a mid-range transmitter and one model-180 sensor placed on the back of the participant’s hand. Due to technical limitations, it was not possible to sample both the position and force data simultaneously. Because the hand position data was only used to determine the timing of the second cue (see Experimental procedure below), at this point the sampling of the trakSTAR was stopped and, after a small delay due to cue presentation and other computer processes (about 400 ms), sampling of the force data began.

The manipulandum was placed on a platform (blue box in Fig. 1b), behind two screens that could switch between an opaque and transparent state (Magic Glass). The top screen provided view of the graspable force sensors (but not the object that was hidden under a paper cover) and the bottom screen of the object in the manipulandum. Due to technical problems with the top screen for three participants in Experiment 1, liquid crystal goggles (PLATO, Translucent Technologies) were used instead that worked similar to the switching screen.

The haptic cues were presented with a haptic device (Phantom 1.5, Sensable), which was controlled with the Haptik Library (de Pascale and Prattichizzo 2007). To make sure the participants did not see the haptic cues, the device was placed in a cardboard box in which a small opening was made where the participant could put in their hand. They rested their hand on a thin cushion on a table with the palm up underneath the device. A magnet was fixed to the arm of the haptic device at which the haptic cues could be attached and easily removed. When the device’s arm was moved in a downward direction, the cue was softly pressed on the palm of the hand and when the arm was moved upward, the cue was released from the participant’s hand.

The force sensors were attached to a NI-USB 6343X (National Instruments) which in turn was connected to a personal computer. The same NI-USB was used to control the switching screens. All systems were controlled and measured with a custom-made script in Matlab (Mathworks).

The objects and cues used in the experiment are shown in Fig. 1a. The objects in Experiment 1 were 3D printed cuboids of different sizes with a width and height of 4 cm and lengths of 4, 7 and 10 cm for the small, medium and large object, respectively. The objects were filled with lead shot and cotton wool to obtain an equal density of 2.5 g/cm^3^for all objects. The resulting weights were approximately 160, 280 and 400 g. Note that the total weight the participants lifted also included the weight of the manipulandum and was 2.8, 3.9 and 5.1 N for the small, medium and large object, respectively. The haptic cues for this experiment were 3D printed plates with the same surface dimensions (4 × 4, 7, or 10 cm), and a height of 0.5 cm. A flat metal ring was taped to the top of the plate to be able to attach it to the magnet on the haptic device.

In Experiment 2, three objects measuring 5 × 5 × 5 cm^3^ were used. The objects were made of balsa wood, cement and aluminium and were referred to the participant as wood, stone and metal. They weighted 64, 195 and 350 g, respectively. When the weight of the manipulandum was included, total weights of 1.8, 3.1 and 4.6 N were obtained. The haptic cues were plates of 5 × 5 × 1 cm^3^, consisting of the same materials. Similar to Experiment 1, a flat metal ring was taped to the top of the plate to attach it to the haptic device.

### 2.3 Experimental procedure

In both experiments, participants received two cues before lifting the object, one before and one after movement initiation. In no-change conditions, the first and second cue were the same. In change conditions, the first cue was incorrect and differed from the second cue. The cues could be presented in three sensory modality conditions: vision, haptic or visuohaptic. Note that although the objects could differ between the first and second cue, the presented modality for both cues always was the same.

The experimental time line is illustrated in Fig. 1c. Participants were seated at a table in front of the two screens. They placed their right hand on the table at the start position (Fig. 1b) and their left hand in the carboard box. Then, participants received a cue indicating which object they would have to grasp. Before the presentation of the cue, the top screen was also turned into a transparent state to indicate that the cue would be presented soon. In addition, the first cue was accompanied by an auditory beep signal to attend the participant of the cue. In the case of a visual cue, the bottom screen became transparent for 1 s, whereas in the case of a haptic cue, the haptic device pressed the plate with a force of 1 N on the left hand of the participant for 1 s. They were allowed to move their fingers to feel the plate, but not to grasp it with their hand. In the visuohaptic condition, the visual and haptic cue were presented simultaneously. There was a small delay between the sending of the command to the haptic device and contact of the haptic device with the hand (on average 140 ms). To make sure the visual and haptic cue were approximately presented at the same time, the visual cue was always delayed with 140 ms as well.

After presentation of the first cue, the top screen became opaque again and the experimenter changed the objects. When the cue was correct (no-change condition), the same object was placed in the manipulandum and the same cue was placed in the haptic device. If the cue was incorrect (change condition), a different object or haptic cue was placed. Next, an audible ‘GO’-signal was played with a computer voice to indicate that the participant could initiate the movement. At the same time, the top screen turned into a transparent state, so the participants could see the graspable surfaces of the manipulandum to guide their movement. When the right hand reached a threshold height (height of 15.2 cm above the table), the second cue was presented. The second cue was presented in the same modality (vision, haptic or visuohaptic) as the first cue and would stay on for the duration of the trial (4 seconds in Experiment 1 and 3 seconds in Experiment 2). The time between the presentation of the second cue and the grasping of the object depended on the movement speed of the participants, but was on average 590 ms (Table 1, time to contact). At the end of the trial, the top screen would turn opaque again and participants replaced the manipulandum on the platform and put their right hand back on the table. The experimenter then replaced the objects and haptic cue for the next trial.

**Table 1.** Results for the time to contact and preloading phase duration. Values indicate mean±standard error and are shown for each modality condition (vision, haptic, visuohaptic), object weight (L=light, H=heavy) and cue change condition (NC=no change, C=change).

|  |  | Vision |  |  |  | Haptic |  |  |  | Visuohaptic |  |  |  |
| --- | --- | --- | --- | --- | --- | --- | --- | --- | --- | --- | --- | --- | --- |
|  |  | L |  | H |  | L |  | H |  | L |  | H |  |
|  | Experiment | NC | C | NC | C | NC | C | NC | C | NC | C | NC | C |
| Time to contact (s) | 1 | 0.55±0.03 | 0.56±0.04 | 0.56±0.03 | 0.55±0.03 | 0.54±0.03 | 0.55±0.03 | 0.54±0.03 | 0.55±0.03 | 0.55±0.03 | 0.55±0.03 | 0.55±0.03 | 0.54±0.04 |
|  | 2 | 0.63±0.04 | 0.63±0.04 | 0.63±0.04 | 0.63±0.04 | 0.63±0.03 | 0.62±0.03 | 0.62±0.04 | 0.62±0.04 | 0.64±0.04 | 0.64±0.04 | 0.64±0.04 | 0.64±0.04 |
| Preloading phase duration (s) | 1 | 0.09±0.01 | 0.08±0.01 | 0.09±0.01 | 0.08±0.01 | 0.08±0.01 | 0.08±0.01 | 0.07±0.01 | 0.07±0.01 | 0.08±0.01 | 0.07±0.01 | 0.07±0.01 | 0.07±0.02 |
|  | 2 | 0.07±0.01 | 0.07±0.01 | 0.08±0.01 | 0.07±0.00 | 0.07±0.00 | 0.07±0.01 | 0.08±0.01 | 0.08±0.01 | 0.07±0.01 | 0.08±0.01 | 0.07±0.01 | 0.09±0.01 |

Participants were instructed to lift the object by placing the thumb and index finger on the force sensors. They moved their hand over the side of the bottom screen to grasp the manipulandum, lift it until the bottom was at the same height as the bottom screen (approximately 5 cm) and hold it steady until the top screen turned opaque again. Then, they replaced the manipulandum on the platform and returned their hand to the start position on the table. It was explained to the participants that they could use the first cue to plan their movement and the correct amount of force. However, if the first cue proved to be incorrect, they should correct their planned movement. They were told not to stop their movement in such a case, but to always move in a fluent motion to grasp and lift the object and move at a comfortable speed.

Before the start of the experiment, participants were familiarized with the haptic cues by pressing each one on their hand with the haptic device. After this, they performed 9 practice trials to familiarize themselves with the procedure and the weights of the different objects. Of these practice trials, the last 2 trials contained a change between the two cues.

### 2.4 Experimental conditions

The no-change trials could consist of the lift of a light, medium or heavy object. These trials will be referred to as LL, MM or HH trials, where L, M and H stand for light, medium and heavy, respectively, and the first letter refers to the first cue and the second to the second cue. The change trials could consist of a light object in the first cue and a heavy object for the second cue (LH trial) or a heavy object in the first cue and a light object for the second cue (HL trial). Only changes between light and heavy objects were analysed because 1) this requires a large difference in motor planning which is easier to measure, and 2) including changes with medium objects would result into a large number of trials which would be demanding for the participants. The medium object was still included in the no-change trials (and a few change trials, see below) to increase task difficulty and make sure that participants would scale their forces more precisely to the weight of each object.

To make sure participants relied on the first cue to plan their movement, there were only change conditions in 1/3 of the trials. For each modality, 16 LL, MM and HH trials were presented (total 144 no-change trials) and 10 LH and HL trials (60 change trials). To ascertain that participants would not expect the medium objects to never change between the first and second cue, 4 trials for each modality were included in which a change with a medium object occurred (random selection of LM, ML, HM or MH trial), which gave another 12 change trials, which were not analysed. Each experiment was divided into two sessions of 108 trials, where each condition was equally divided across the sessions. Within each session, the conditions were randomly presented, but no more than 2 change trials were presented in a row. Furthermore, the first 9 trials always consisted of the 9 no-change trials for each modality and each weight in a random order, to make sure each condition and object was experienced at the start of the experiment.

### 2.5 Data analysis

The position data was only used to determine the timing of the second cue and was not further analysed. The force data was filtered with a 2^nd^ order bidirectional lowpass Butterworth filter with a cut-off frequency of 15 Hz. The load force (LF) and grip force (GF) were defined as the sum of the vertical forces of both sensors and the mean of the horizontal forces, respectively. The force rates (LFR and GFR) were calculated as the first derivative of LF and GF with respect to time. The onset of LF and GF were determined when the forces reached a threshold of 0.1 N. This value was chosen to exclude small variations due to noise when bumping against the manipulandum when grasping the force sensors.

The parameters of interest were the first peak of the force rates (peakGFR1 and peakLFR1). These parameters reflect the force scaling, where especially the first peaks indicate the planned force before feedback processes are included (Johansson and Westling 1988). The first peak of force rates was the first peak that occurred between GF onset and just after lift-off (LF overcomes object weight). To exclude small peaks that are due to noise, the first peak had to minimal 30% of the maximum peak within this time frame (van Polanen et al. 2020).

Even though participants were instructed to continue their movement and not stop to wait for the second cue, it is possible that they changed their behaviour when the second cue was different from the first. To investigate whether they moved slower at the final phase of their movement, the time to contact and the preloading phase were calculated. The time to contact was the time between the second cue and object contact (GF onset) and was the final part of the reaching phase. The preloading phase was the time between GF onset and LF onset and reflects the duration between grasping the object and the start of force increments.

For all parameters, values were averaged for each condition (3 modalities, 3 weights and 3 cues). Trials in which participants lifted the object incorrectly or before the go-signal (Exp 1: 16 trials 1.5%, Exp 2: 6 trials, 0.2%), the wrong object was placed (Exp 1: 2 trials 0.1%, Exp 2: 1 trial <0.1%) or a technical problem occurred (Exp 1: 0 trials 0%, Exp 2: 3 trials 0.1%) were removed from the analysis. Because the force data collection was started after participants reached a threshold height, sometimes participants that moved quickly already grasped the object before the start of the force data collection. When at the start of the measurement the load force was above 0.1 N and/or the grip force was above 0.5 N, trials were removed as well (Exp 1: 28 trials, 0.9%, Exp 2: 1 trial, <0.1%). These thresholds were chosen because then the first peaks of the force rates could still be determined. For the time to contact and the preloading phase, the correct determination of the GF onset (GF>0.1 N) was important as well. Therefore, for these variables, trials that started with values of GF above 0.1 were removed as well (Exp 1: 50 trials, 1.5%, Exp 2: 2 trials, 0.1%). In addition, for these two parameters, there were sometimes large outliers. Therefore, an outlier analysis was performed and values that deviated more than 3 standard deviations from the mean were removed for the time to contact and the preloading phase (Exp 1: 22 and 50 trials; Exp 2: 27 and 46 trials for the time to contact and preloading phase duration, respectively).

### 2.6 Statistics

All analyses were performed separately for both experiments. First, it was determined whether participants corrected their force planning towards the three object weights. This analysis was performed on the conditions where no change in object took place (LL, MM and HH) as a baseline comparison. A 3 (object weight: light, medium or heavy) × 3 (modality: vision, haptic or visuohaptic) repeated measures analysis of variance (ANOVA) was performed on the peakGFR1 and peakLFR1.

To test whether there were differences between conditions when the object was changed between the first and second cue (change conditions: LH and HL) and when it remained the same (no-change conditions: LL and HH), a 2 (cue change: change or no-change) × 2 (weight: light or heavy) × 3 (modality: vision, haptic or visuohaptic) repeated measures ANOVA was conducted on the peakGFR1, peakLFR1, preloading phase duration and time to contact. Since there were only a few trials in which the object changed from or to a medium weight, this object was not included in this analysis.

If the sphericity assumption was violated (Maughly’s test), a Greenhouse-Geisser correction was used. Post-hoc tests were performed with paired samples t-tests with a Bonferroni correction. A p-value <0.05 was considered significant.

### 2.7 Force scaling effects and multisensory integration

Optimal multisensory integration can be represented with maximum-likelihood integration (Ernst and Banks 2002). If sensory estimates are modelled as Gaussian distributions with a variance σ^2^, the maximum-likelihood rule states that unimodal estimates are integrated according to their variance. That is, the multimodal variance can be calculated from the unimodal variances and would have a lower variance than the unimodal ones: 

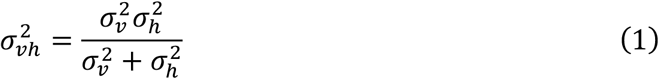

To obtain optimal integration of vision and haptics, the modalities are combined and weighted according to their reliability. The larger the variance, the smaller this reliability and the lower the weighting given to this modality. The weighting is calculated using the variance for each modality and the sum of the variances of all modalities:

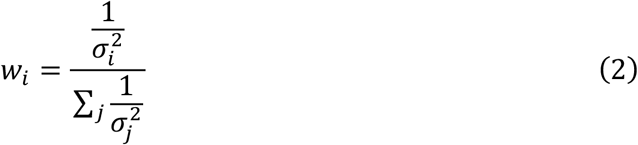

The weightings of vision (*w*_*v*_) and haptics (*w*_*h*_) will sum up to 1. In perception research, the variance is usually estimated by measuring the discrimination threshold. In the present experiment, this threshold would be difficult to measure, since only a few object weights were used. However, the difference in force scaling between the light and the heavy object could be seen as a measure of discrimination. In other words, a larger difference between the light and heavy object would indicate a more precise scaling and a more precise discrimination. In this case, the force scaling difference (Δ*F*) would be inversely related to the variance:

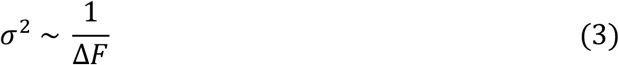

For calculation of Δ*F*, the peakGFR1 and the peakLFR1 were used in separate analyses. It was calculated for no-change (HH-LL) and change conditions (LH-HL) and for each modality condition. When participants scale their forces differently for the light and heavy object, ΔF should be significantly different from zero. This was determined by using one sample t-tests.

The average over all participants of the force discrimination values for the haptic and visual modality were used to predict the difference in the visuohaptic condition by combining equation (1) and (3) and compare this to the measured difference to determine whether the unimodal signals were integrated according to the optimal integration principle. The predicted values were compared to the measured differences in the visuohaptic condition with one sample t-tests, separately for change and no-change conditions, and peakGFR1 and peakLFR1.

The force scaling difference was also used to calculate the weight of each modality (equation (3)), by replacing σ^2^ with Δ*F*, using equation (2). The weights were determined for both the change and no-change conditions, based on the peakGFR1 and on the peakLFR1.

## 3 RESULTS

In this study, two experiments were performed on the corrections of force scaling in object lifting based on (multi)sensory cues. All data used for analyses and figures can be found at https://osf.io/d2afe/.

### 3.1 Force scaling for three object weights

As a first step, it was examined whether participants scaled their forces to the different weights. Main effects of weight were found for all force parameters in both experiments (Fig. 2).

**Fig. 2.**
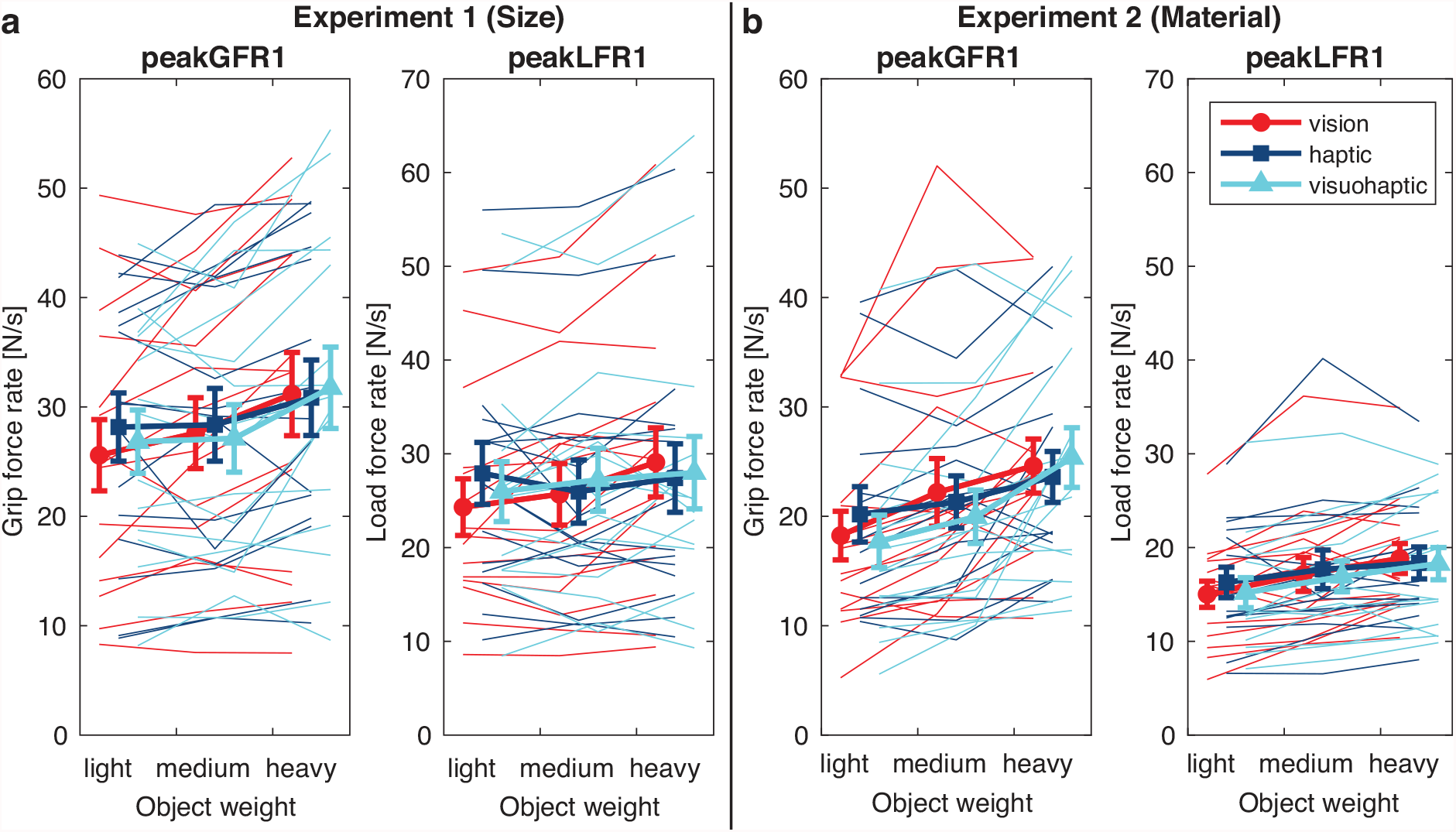
Force scaling for three weights (light, medium and heavy) in the no-change conditions for the first peak of grip force rate (peakGFR1) and first peak of load force rate (peakLFR1). for Experiment 1 (a) and Experiment 2 (b). Markers indicate the three modality conditions (vision, haptic and visuohaptic) with standard errors. Thin lines represent individual participants. Note that a main effect of weight was found for all parameters in both experiments

#### 3.1.1 Experiment 1: forces were scaled to size cues

In Experiment 1, the 3 (weight) × 3 (modality) ANOVA on peakGFR1 only showed a main effect of weight (F(2,28)=19.0, p<.001, *η*_p_^2^=0.58). Post-hoc tests indicated that the heavy weight (31.3±3.6) was lifted with higher grip forces than the light weight (26.9±3.0, p<0.001) and the medium weight (27.7±3.1, p=0.002), but the light and medium weight did not differ (p=0.385).

For the peakLFR1, also a main effect of weight was found (F(2,28)=4.9, p=0.015, *η*_p_^2^=0.26). However, in the post-hoc tests, no significant differences were found between the light (26.1±3.1 N/s, p=0.049), medium (26.3±3.1 N/s) and heavy (28.2±3.7 N/s) weight. However, also an interaction between weight × modality was seen (F(4,56)=4.1, p=0.006 *η*_p_^2^=0.23). Post-hoc tests indicated that a difference between light and heavy objects was found for the vision condition (p=0.018), but not for the haptic or visuohaptic condition (p>0.10). No other differences were found between the weights or modalities for peakLFR1.

#### 3.1.2 Experiment 2: forces were scaled to material cues

For the second experiment, a main effect of weight was found for peakGFR1 (F(1.3,18.6)=14.6, p<.001, *η*_p_^2^=0.51). The light object was lifted with a lower peak force rate (18.7±2.3 N/s) compared to the medium object (p=0.003, 21.2±2.6 N/s) and the heavy object (p=0.001, 24.5±2.4 N/s), while the latter two did not differ significantly (p=0.059). However, an interaction between weight and modality (F(4,56)=3.0, p=0.026, *η*_p_^2^=0.18) indicated that the difference between a light and heavy object was only seen in the vision (p=0.12) and visuohaptic (p=0.036) modality conditions, but not the haptic one (p=0.198). No other post-hoc tests were significant.

For peakLFR1 in Experiment 2, also a main effect of weight was found (F(2,28)=20.4, p<.001, *η*_p_^2^=0.59). The light object was lifted with a lower force rate (15.5±1.5 N/s) compared to the medium (p=0.007, 17.2±1.8 N/s) and the heavy object (p<0.001, 18.5±1.6 N/s), whereas the latter two did not differ significantly (p=0.121).

In sum, in both experiments the participants scaled their forces according to the weight of the object, where force rates increased with object weight. No differences between the modalities were found, although the difference between the weights did sometimes depend on the modality. In general, the force scaling appeared to be weakest when the cue was presented only in the haptic modality.

### 3.2 Force scaling can be corrected with a change in object weight

The previous analysis showed that participants scaled their forces towards the object weight when the first cue was correct and the object was not changed. To see whether this behaviour was different when the cue was incorrect and force scaling needed to be adapted, the change and no-change conditions were compared. As can be seen in Fig. 3, there were no differences between these conditions in all modality conditions.

**Fig. 3.**
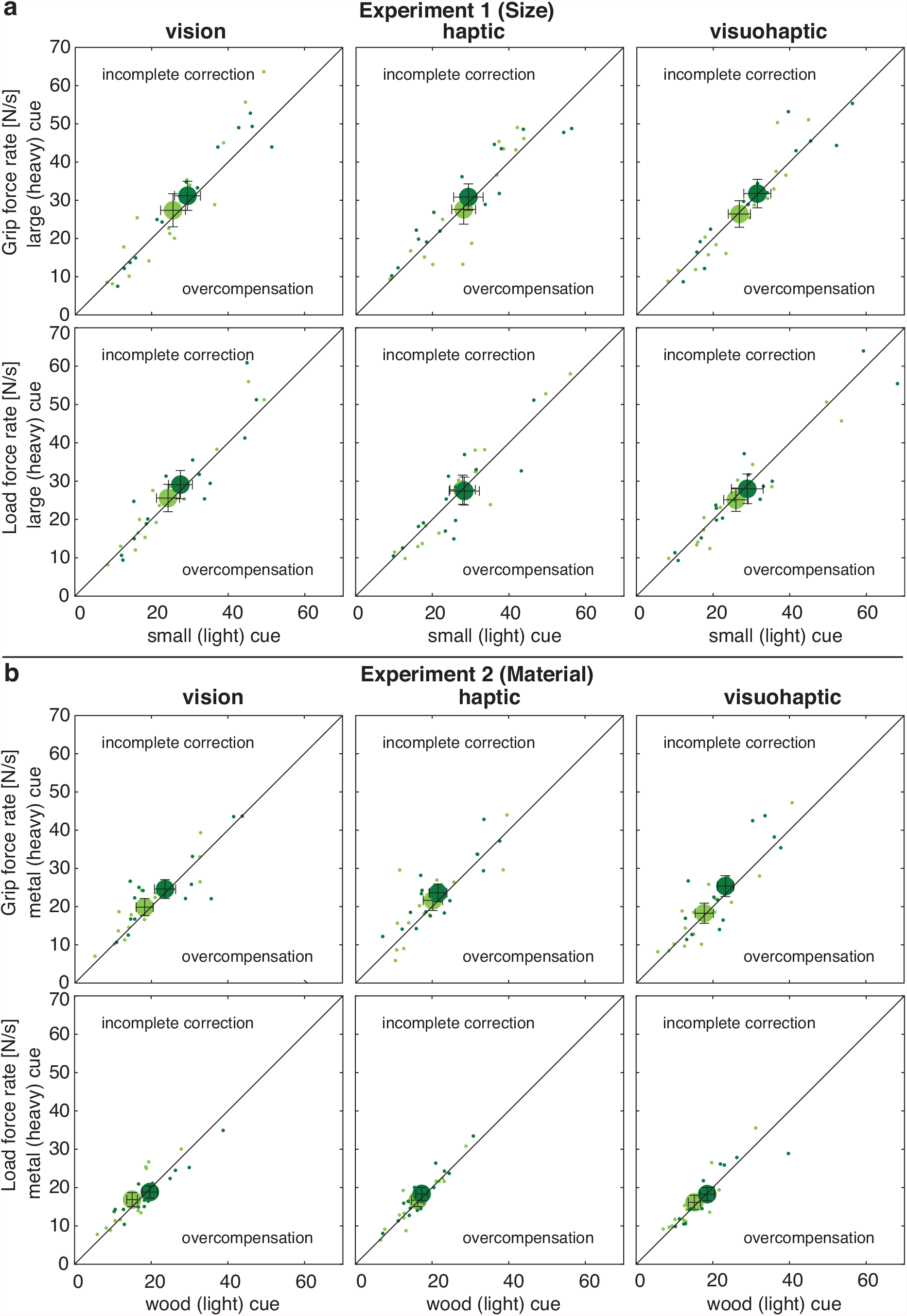
Differences between change and no-change conditions for Experiment 1 (a) and Experiment 2 (b) in the three modality conditions (vision, haptic and visuohaptic), for the first peak of grip force rate (peakGFR1) and first peak of load force rate (peakLFR1). Light and dark green symbols represent light and heavy objects, respectively. The lifted object is plotted by comparing the first cue when it was small (light, x-axis) or large (heavy, y-axis). If the object is lifted similarly, irrespective of the first cue, values will fall on the diagonal black line. If the value is different from the black line, there is either incomplete correction (force is scaled towards the first cue and less to the object weight) or overcompensation (force is scaled towards the object weight, but more than in the no-change condition). Small dots represent values for individual participants. Error bars indicate standard errors. Note that no significant differences were found between change and no-change conditions

#### 3.2.1 Experiment 1: forces can be corrected based on size cues

In Experiment 1, similar to the previous analysis, it was found that the force parameters depended on object weight, but not on cue change. For the peakGFR1, the 2 (cue change) × 2 (weight) × 3 (modality) ANOVA revealed only an effect of weight (F(1,14)=18.4, p<.001, *η*_p_^2^=0.57), with a lower peak for light compared to heavy lifts. For the peakLFR1, the same effect of weight was seen (F(1,14)=11.0, p=0.005, *η*_p_^2^=0.44). For both parameters, no effect of cue change was observed.

### 3.2.2 Experiment 2: forces can be corrected based on material cues

In the second experiment, similar results were found as in Experiment 1. There was an effect of weight for the peakGFR1 (F(1,14)=15.8, p=0.001, *η*_p_^2^=0.53): the force rate peak was lower for the light compared to the heavy object. However, a weight × modality interaction (F(2,28)=4.3, p=0.024, *η*_p_^2^=0.23) revealed that this difference was only found in the visuohaptic modality condition (p<0.018). No significant differences between the modality conditions were found.

For the peakLFR1 in Experiment 2, also a difference between the weights was found (F(1,14)=80.6, p<.001, *η*_p_^2^=0.85) where the force rate peak was lower for the light than the heavy object. There was also an interaction between weight × modality for the peakLFR1 (F(2,28)=3.5, p=0.042, *η*_p_^2^=0.20). Post-hoc tests showed that the difference between light and heavy weights was only seen in the vision (p<0.001) and the visuohaptic (0.002) modality conditions. No differences between the modalities were found.

In sum, the differences between the object weights as found in the first analysis were again found in this analysis. Importantly, no effects of cue change were found, indicating that the force scaling was not different between conditions in which the object was correctly or incorrectly indicated with the first cue. This was the case for both experiments, whether objects were indicated with size or material cues, and for all modality conditions, whether cues were indicated visually, haptically or both.

### 3.3 Multisensory integration of visual and haptic information

The differences in force scaling between heavy and light objects was used as a measure of discrimination, and these differences are illustrated in Fig. 4. The larger this difference, the higher discrimination performance. Significant differences from zero are indicated with blue asterisks. The values seem larger in the no-change condition (purple bars) than in the no-change conditions (cyan bars), but a repeated measures 2 (cue change) × 3 (modality) ANOVA revealed no main effects or interactions with the factor cue change, indicating no significant difference between conditions with or without a cue change.

**Fig. 4.**
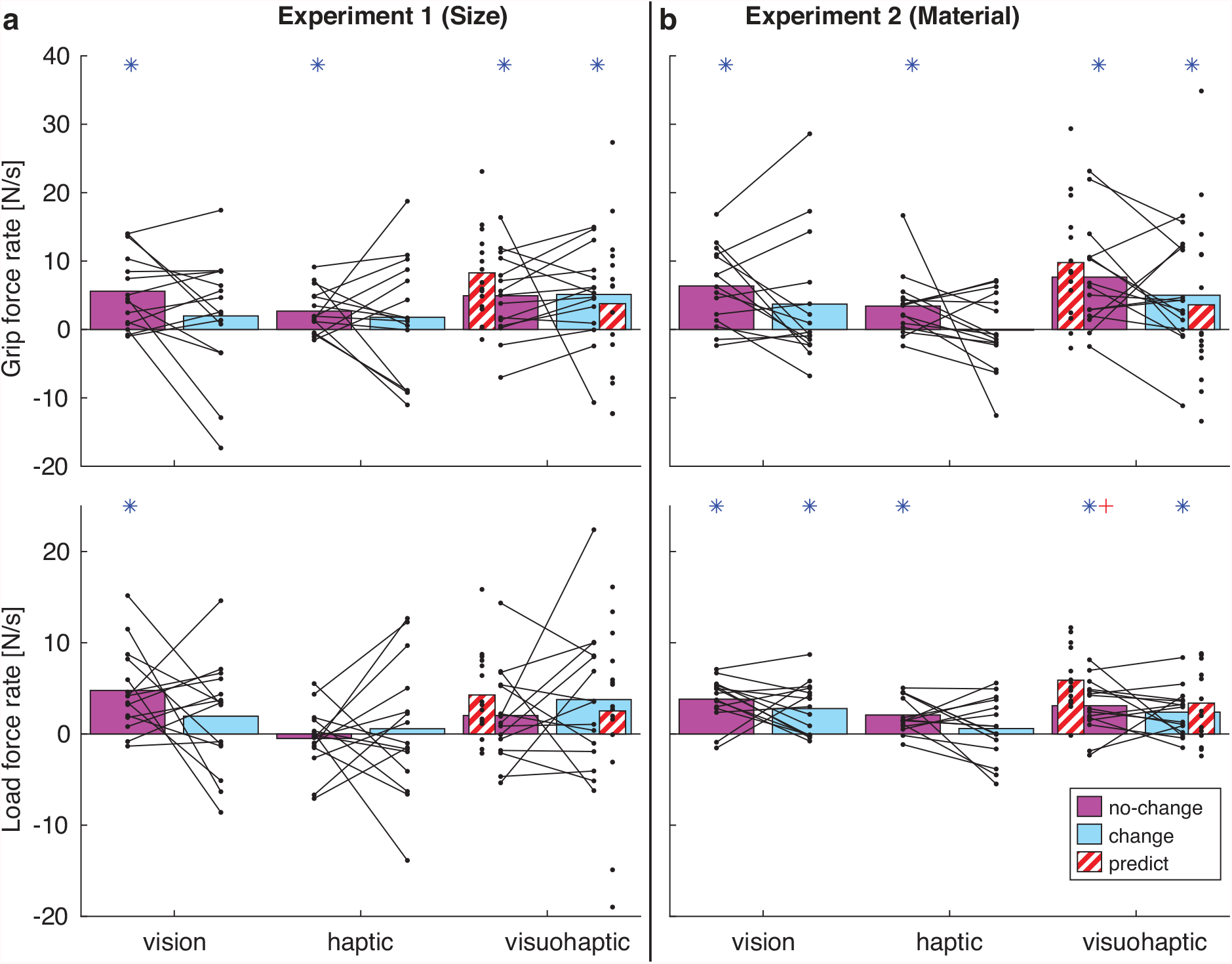
Force scaling differences (heavy – light) for no-change and change conditions in each modality, for Experiment 1 (a) and Experiment 2 (b). Top and bottom panels show the force scaling based on first peaks of grip force rate and load force rates, respectively. Larger bars represent higher discrimination between light and heavy objects. Red striped bars are the predicted force scaling for the visuohaptic condition, based on the vision and haptic conditions. Black dots and lines represent individual participants. * significant difference from zero (one-sample t-test). + significant difference between predicted and measured visuohaptic force scaling differences

Based on the force differences in the vision and haptic conditions, a predicted force scaling difference was calculated for the visuohaptic condition to see whether the two modalities were integrated in an optimal fashion. The predicted force differences (red striped bars in Fig. 4) were quite accurate, as they did not differ from the measured differences in the visuohaptic condition (all p>0.13), except in Experiment 2 for peakLFR1 in the no-change condition (p=0.003, red plus sign in Fig. 4).

The weightings for the no-change conditions indicated that visual information was weighted higher (>0.5) than haptics in both experiments (Exp 1 peakGFR1: *w*_*v*_=0.68, peakLFR1: *w*_*v*_=1.12; Exp 2 peakGFR1: *w*_*v*_=0.65, peakLFR1: *w*_*v*_=0.65). In the change conditions, the weighting for vision decreased in Experiment 1 (peakGFR1: *w*_*v*_=0.53, peakLFR1: *w*_*v*_=0.77), whereas the weighting for vision increased for Experiment 2 (peakGFR1: *w*_*v*_=1.03, peakLFR1: *w*_*v*_=0.82). Note that values of *w*_*v*_ above 1 were found if the force difference for haptics was below zero, indicating that forces were scaled higher for light objects compared to heavy, which results in a negative force difference and a negative weighting for haptics.

To check whether there were significant differences between the conditions in how visual and haptic information was integrated, the weightings were also calculated for the individual participants. However, since many participants had a force difference below zero, this could lead to negative weighting values, which would be difficult to interpret. To avoid this problem, all force differences were increased by 20 N/s, to make sure all values would be above zero. The downside of this method is that the relative differences between the conditions would become smaller, which would also lead to smaller differences in the weightings for vision and haptics. However, it would be possible to test whether the weightings would differ significantly between the conditions. Paired t-tests showed that *w*_*v*_ did not differ between change and no-change conditions, neither for peakLFR1 or peakGFR1 and in neither experiment (all p>0.32).

To test whether the weighting was larger for vision or haptics, one sample t-tests were used to determine a significant difference from 0.5. A larger weighting for vision (*w*_*v*_>0.5) was found for peakLFR1 in the no-change conditions in Experiment 1 (p=0.006) and Experiment 2 (p=0.046). However, for the change conditions for peakLFR1, no significant difference from 0.5 was found, nor for peakGFR1 in any condition, suggesting an equal weighting of vision and haptics.

In sum, the visuohaptic discrimination value could be predicted from the visual and haptic values in most conditions, which suggests that both sources of information were integrated in an optimal way. Both visual and haptic information were used in the visuohaptic condition, with a higher weighting for vision in some conditions. However, there were no significant differences between change and no-change conditions.

### 3.4 Reach and grasp parameters are not largely affected by object change

To test whether reach and grasp behaviour was altered with an object change after the first cue, the time to contact and the preloading phase duration were calculated (Table 1). For the time to contact, no significant effects of cue change, weight or modality were observed in neither experiments, indicating no difference in reaching behaviour. For the preloading phase duration in Experiment 1, there was an effect of weight (F(1,14)=10.0, p=0.007, *η*_p_^2^=0.42), where the duration was longer for light compared to heavy objects. In Experiment 2, also an effect of weight was found (F(1,14)=9.3, p=0.009, *η*_p_^2^=0.40), but with a longer duration for heavy compared to light objects. Furthermore, there was a significant interaction between cue change and modality (F(2,28)=8.2, p=0.002, *η*_p_^2^=0.37). In the visuohaptic condition, the duration was shorter when there was no change (0.069±0.005 s) than when there was an object change (0.083±0.006 s; p=0.002). There were no differences in the other modality conditions.

## 4 DISCUSSION

The main aim of the experiment was to investigate whether fast corrections could be made to a prepared motor plan in response to a change in sensory information about object properties, even after the initiation of the movement. In the experiments, three different weights were used that were indicated with three different sizes (Experiment 1) or materials (Experiment 2). In both experiments, participants were able to use the information of the cues to anticipatory plan their forces to the weight of the objects. That is, the force scaling as indicated by the first peaks in grip and load force rates, increased with the weight of the lifted object. During the reaching movement, the cue could change to indicate a different weight and participants had to quickly adjust their planned forces during reaching in order to lift the object skilfully. Previous studies showed that changes in visual size information (Brouwer et al. 2006) and colour cues (van Nuenen et al. 2012) after the movement was initiated could be used to adjust the planned fingertip forces. The current study extends these findings by showing that 1) in addition to size and colour information, also material information can be used to correct planned forces 2) in addition to visual information, sensory cues presented haptically or visuohaptically can also be used to correct the motor plan.

Participants had on average 590 ms (time to contact, range 370-940 ms) before they contacted the object and 76 ms (preloading phase, range 32-208 ms) to process the cue change and adjust their motor plan. This time frame of 666 ms was on average enough to enable participants to correct their initial motor plan and scale their forces to the new weight. The present study cannot state what minimum time interval is needed in order to make online corrections, since participants could choose their own comfortable movement speed and the timing of the second cue was not varied. With average reach times of about 900 ms and an object change around 470 ms, the study of Brouwer et al. (2006) suggests that even shorter time windows (430 ms) are possible.

In case of a cue change and the need for corrections, participants did not slow down their reaching movement to allow them more time for changing the plan. In one condition, there was a small increase (15 ms on average) in preloading phase, but it does not seem likely that this would completely cover the time needed for corrections. This suggests that participants were able to correct their motor plan online during the movement. In the original dual-stream theory, online corrections are processed by the dorsal stream (Goodale and Milner 1992), which would not have quick access to memorized information, such as an association between a perceptual cue and object weight. The present results are not in line with that view and suggest that information is quickly available. Two possible explanations can be proposed. First, the dorsal stream could have processed some cue information linked with object weight and immediately induced the required corrections to the motor plan without need of the ventral stream. For instance, previous research suggests that size information is processed in the intraparietal area (Murata et al. 2000; Chouinard et al. 2009; Monaco et al. 2015), which is part of the dorsal stream. However, Experiment 2 showed that corrections could also be based on material information. Since texture information, indicative of material, is known to be processed in the ventral stream (James et al. 2007; Cant et al. 2009; Kim et al. 2019), it is unlikely that the processing of the cues information about the object’s weight and the implementation of corrections could be solely attributed to the dorsal stream. A second, more likely, explanation is that the dorsal stream receives input from the ventral stream, that are known to be connected from human and primate studies (Distler et al. 1993; Borra et al. 2008; Takemura et al. 2016; Budisavljevic et al. 2018). The present study shows that information between these streams can be shared at a relatively short time scale during ongoing movements. Future work should study the brain areas involved in processing information for online corrections. For instance, it is possible that one central area collects different sources of information to implement corrections.

A second aim of this study was to investigate how multisensory information was integrated in the online corrections of lifting movements. The predicted force discrimination values from the maximum-likelihood estimation based on the vision and haptic conditions were similar to the actual discrimination values in the visuohaptic condition. This suggests that information from vision and haptics was integrated in an optimal fashion, i.e. combining the information from both sources in an optimal way based on their reliability. Nevertheless, in one condition the predicted value differed from the measured value. Furthermore, despite the availability of multiple sources of information, the visuohaptic condition did not outperform the visual and haptic conditions, which would be expected with optimal integration. Therefore, it seems that both visual and haptic information is used in the visuohaptic condition, but it is unclear whether information is integrated optimally.

Notably, force scaling seemed to be less tuned to the object weight when information was presented haptically. Even though participants remarked that they could distinguish the haptic cues, it might be possible that it was more difficult to perceive the haptic cue compared to the visual cue. The haptic cue was presented to the non-grasping left hand at a different location than the object, which could have made the association between the cue and the object less strong. Although previous literature showed that weight information from one hand can be transferred to the other hand for force scaling (Gordon et al. 1994; Chang et al. 2008), indicating that information is available for the other hand, it is reasonable to assume that it takes longer to process information that is presented to the other hand. Another explanation could be that a short movement with the fingers was needed to perceive the haptic cue, which takes time. Especially for the material cues, both the wooden (light) and metal (heavy) object were quite smooth, which might have made them more difficult to distinguish haptically, whereas they were very different in colour and could be quickly discriminated visually.

The decreased force discrimination with haptic information suggests that visual information has a larger contribution compared to haptic information when visual and haptic information are combined in the visuohaptic condition. Indeed, the weights found for vision were larger compared to haptics. This corresponds to the grasping study of (Pettypiece et al. 2010), that found that there was more relied on visual information about object size. A previous object lifting study found a slightly larger weighting for haptics compared to vision (van Polanen et al. 2019), but here the sensory information referred to position of the fingers, not the object properties. It must be also noted that the variability in discrimination performance between individuals is large. Perhaps the optimal weighting between vision and haptics was also different for different participants, for example, depending on how well they could haptically discriminate the objects. Finally, the weighting of haptic and visual information did not seem to depend on whether corrections needed to be made, or the type of visual information (size or material). Possibly, the weighting of haptic and visual information depends on the specific task or motor process that needs to be controlled, and the source of the information (e.g. from the object or the hand), but not the type of information.

In conclusion, the present findings show that a motor plan of a lifting movement can be quickly corrected after the movement has been initiated based on cued information that is linked to a stored object weight. Cued information from different object properties (size and material) and presented towards different modalities (vision and haptics) can be used to make such corrections. In addition, visual and haptic information is integrated when multisensory information is available. These findings provide insights into the information that can be shared between brain areas for the online control of hand-object interactions.

## CONFLICT OF INTEREST

The author declares no competing financial or non-financial interests.

## DATA AVAILABILITY

All data used for analyses and figures can be found at https://osf.io/d2afe/.

## ACKNOWLEDGEMENTS

The author would like to thank Silke Thys for her help in data collection for Experiment 2 and Guy Rens for providing comments to an earlier version of the manuscript.

## Notes

https://osf.io/d2afe

### Competing Interest Statement

The authors have declared no competing interest.

## REFERENCES

Betti S, Castiello U, Begliomini C (2021) Reach-to-Grasp: A Multisensory Experience. Frontiers in Psychology 12:213

Borra E, Belmalih A, Calzavara R, Gerbella M, Murata A, Rozzi S, Luppino G (2008) Cortical connections of the macaque anterior intraparietal (AIP) area. Cereb Cortex 18:1094–1111 doi: 10.1093/cercor/bhm146

Brouwer AM, Georgiou I, Glover S, Castiello U (2006) Adjusting reach to lift movements to sudden visible changes in target’s weight. Exp Brain Res 173:629–636 doi: 10.1007/s00221-006-0406-x

Buckingham G, Cant JS, Goodale MA (2009) Living in A Material World: How Visual Cues to Material Properties Affect the Way That We Lift Objects and Perceive Their Weight. Journal of Neurophysiology 102:3111–3118 doi: 10.1152/jn.00515.2009

Budisavljevic S, Dell’Acqua F, Castiello U (2018) Cross-talk connections underlying dorsal and ventral stream integration during hand actions. Cortex 103:224–239 doi: 10.1016/j.cortex.2018.02.016

Camponogara I, Volcic R (2019a) Grasping adjustments to haptic, visual, and visuo-haptic object perturbations are contingent on the sensory modality. J Neurophysiol 122:2614–2620 doi: 10.1152/jn.00452.2019

Camponogara I, Volcic R (2019b) Grasping movements toward seen and handheld objects. Sci Rep 9:3665 doi: 10.1038/s41598-018-38277-w

Cant JS, Arnott SR, Goodale MA (2009) fMR-adaptation reveals separate processing regions for the perception of form and texture in the human ventral stream. Experimental Brain Research 192:391–405 doi: 10.1007/s00221-008-1573-8

Chang EC, Flanagan JR, Goodale MA (2008) The intermanual transfer of anticipatory force control in precision grip lifting is not influenced by the perception of weight. Experimental brain research 185:319–329 doi: 10.1007/s00221-007-1156-0

Chouinard PA, Large ME, Chang EC, Goodale MA (2009) Dissociable neural mechanisms for determining the perceived heaviness of objects and the predicted weight of objects during lifting: an fMRI investigation of the size-weight illusion. Neuroimage 44:200–212 doi: 10.1016/j.neuroimage.2008.08.023

Cloutman LL (2013) Interaction between dorsal and ventral processing streams: where, when and how? Brain Lang 127:251–263 doi: 10.1016/j.bandl.2012.08.003

Crevecoeur F, Munoz DP, Scott SH (2016) Dynamic Multisensory Integration: Somatosensory Speed Trumps Visual Accuracy during Feedback Control. The Journal of Neuroscience 36:8598–8611 doi: 10.1523/JNEUROSCI.0184-16.2016

de Pascale M, Prattichizzo D (2007) The Haptik Library: A Component Based Architecture for Uniform Access to Haptic Devices. IEEE Robotics & Automation Magazine 14:64–75

Dijkerman HC, de Haan EH (2007) Somatosensory processes subserving perception and action. Behav Brain Sci 30:189–201 doi: 10.1017/S0140525X07001392

Distler C, Boussaoud D, Desimone R, Ungerleider LG (1993) Cortical connections of inferior temporal area TEO in macaque monkeys. J Comp Neurol 334:125–150 doi: 10.1002/cne.903340111

Ernst MO, Banks MS (2002) Humans integrate visual and haptic information in a statistically optimal fashion. Nature 415:429–433 doi: 10.1038/415429a

Gallivan JP, Cant JS, Goodale MA, Flanagan JR (2014) Representation of object weight in human ventral visual cortex. Curr Biol 24:1866–1873 doi: 10.1016/j.cub.2014.06.046

Goodale MA, Milner AD (1992) Seperate visual pathways for perception and action. Trends Neurosci 15:20–25

Gordon AM, Forssberg H, Iwasaki N (1994) Formation and lateralization of internal representations underlying motor commands during precision grip. Neuropsychologia 32:555–568

Gordon AM, Forssberg H, Johansson RS, Westling G (1991a) The integration of haptically acquired size information in the programming of precision grip. Exp Brain Res 83:483–488

Gordon AM, Forssberg H, Johansson RS, Westling G (1991b) Visual size cues in the programming of manipulative forces during precision grip. Exp Brain Res 83:477–482

James TW, Kim S, Fisher JS (2007) The neural basis of haptic object processing. Canadian Journal of Experimental Psychology/Revue canadienne de psychologie expérimentale 61:219–229 doi: 10.1037/cjep2007023

Johansson RS, Flanagan JR (2009) Coding and use of tactile signals from the fingertips in object manipulation tasks. Nat Rev Neurosci 10:345–359 doi: 10.1038/nrn2621

Johansson RS, Westling G (1988) Coordinated isometric muscle commands adequately and erroneously programmed for the weight during lifting task with precision grip. Exp Brain Res 71:59–71

Kim T, Bair W, Pasupathy A (2019) Neural coding for shape and texture in macaque area V4. The Journal of Neuroscience 39:3073–3018 doi: 10.1523/JNEUROSCI.3073-18.2019

Milner AD (2017) How do the two visual streams interact with each other? Experimental Brain Research 0:0–0 doi: 10.1007/s00221-017-4917-4

Milner AD, Goodale MA (2008) Two visual systems re-viewed. Neuropsychologia 46:774–785 doi: 10.1016/j.neuropsychologia.2007.10.005

Monaco S, Sedda A, Cavina-Pratesi C, Culham JC (2015) Neural correlates of object size and object location during grasping actions. Eur J Neurosci 41:454–465 doi: 10.1111/ejn.12786

Murata A, Gallese V, Luppino G, Kaseda M, Sakata H (2000) Selectivity for the shape, size, and orientation of objects for grasping in neurons of monkey parietal area AIP. J Neurophysiol 83:2580–2601 doi: 10.1152/jn.2000.83.5.2580

Oldfield RC (1971) The assessment and analysis of handedness: the Edinburgh inventory. Neuropsychologia 9:97–113

Pettypiece CE, Goodale Ma, Culham JC (2010) Integration of haptic and visual size cues in perception and action revealed through cross-modal conflict. Experimental brain research 201:863–873 doi: 10.1007/s00221-009-2101-1

Takemura H, Rokem A, Winawer J, Yeatman JD, Wandell BA, Pestilli F (2016) A Major Human White Matter Pathway Between Dorsal and Ventral Visual Cortex. Cerebral Cortex 26:2205–2214 doi: 10.1093/cercor/bhv064

van Nuenen Bfl, Kuhtz-Buschbeck J, Schulz C, Bloem BR, Siebner HR (2012) Weight-specific anticipatory coding of grip force in human dorsal premotor cortex. J Neurosci 32:5272–5283 doi: 10.1523/JNEUROSCI.5673-11.2012

van Polanen V, Davare M (2015) Interactions between dorsal and ventral streams for controlling skilled grasp. Neuropsychologia 79:186–191 doi: 10.1016/j.neuropsychologia.2015.07.010

van Polanen V, Rens G, Davare M (2020) The role of the anterior intraparietal sulcus and the lateral occipital cortex in fingertip force scaling and weight perception during object lifting. J Neurophysiol 124:557–573 doi: 10.1152/jn.00771.2019

van Polanen V, Tibold R, Nuruki A, Davare M (2019) Visual delay affects force scaling and weight perception during object lifting in virtual reality. J Neurophysiol 121:1398–1409 doi: 10.1152/jn.00396.2018

